# Model-based assessment of race, sex, and electrode montage in ECT

**DOI:** 10.64898/2026.08.20.745969

**Authors:** Niranjan Khadka, Yu Huang, Zhi-De Deng, Dennis Q. Truong, Ganesan Venkatasubramanian, Yiheng Tu, Weiwei Ma, Christopher C Abbott, Abhishek Datta

## Abstract

**Objective:** This computational modeling study quantified the influence of sex and race-related cranial anatomy on predicted brain-wide current flow during electroconvulsive therapy (ECT) across conventional (bifrontal (BF), bitemporal/bilateral (BL), right unilateral (RUL)) and experimental (focal electrically administered seizure therapy (FEAST) and frontomedial (FM)) electrode montages. The objective was to determine whether race-associated variability meaningfully contributes to differences in ECT stimulation metrics across montages.

**Methods:** Finite element head models of Chinese, Black, and Caucasian subjects were developed using high-resolution magnetic resonance imaging and analyzed using the Realistic vOlumetric-Approach-based Stimulator for Transcranial electric stimulation (ROAST) pipeline (N = 150 total; n = 50 per cohort, comprising 25 M and 25 F, age range: 20-30 years). Five ECT montages were simulated under a constant-current condition (900 mA). Stimulation strength (*`E`*_brain_/*E**_th_*) was quantified as 90^th^ percentile of brain-wide E-field magnitude (*`E`_brain_*) relative to neuronal activation threshold (*`E`_th_*= 0.25 V/cm) quantified stimulation strength. Overall focality was evaluated as a percentage of brain volume stimulated above the neural activation threshold (*`E`_brain_ `≥ E`_th_*), while laterality was quantified as the median right-to-left hemispheric E-field magnitude ratio. The effects of race, sex, and montage on stimulation strength, focality, and hemispheric laterality were statistically analyzed.

**Results:** Substantial race- and sex-related differences observed in cranial anatomy resulted in systematic variation in predicted ECT-induced E-field intensity. Brain-wide E-field magnitude varied by both race and montage, with the largest fields generally observed in Caucasian head models and during BL stimulation. Montage exerted the strongest effect on stimulation strength (*`E`_brain_*/*E**_th_*) with BL and FEAST producing the highest stimulation strengths, followed by RUL and FM, while BF produced the lowest. Caucasian subjects generally predicted higher stimulation strengths than Black and Chinese subjects, whereas females predicted modestly higher stimulation strengths than males. Laterality was primarily determined by montage, with FEAST producing the greatest hemispheric asymmetry, followed by RUL. Chinese subjects demonstrated higher laterality ratios than both Black and Caucasian subjects. BL, RUL, and FEAST stimulated substantially larger brain volumes above neural activation threshold (less focal stimulation) than BF. Lower focality was observed in Caucasian subjects relative to Black and Chinese subjects, and in females relative to males.

**Conclusions:** Electrode montage was the primary determinant of predicted ECT stimulation strength, focality, and laterality. Race-related anatomical differences and, to a lesser extent, sex-related differences systematically altered stimulation patterns, supporting consideration of individualized anatomy in ECT dosing and treatment optimization.

## 1. Introduction

Electroconvulsive therapy (ECT), first introduced in the 1930s, remains one of the most effective treatment modalities in psychiatry^1,2^, particularly for severe and treatment-resistant major depressive disorder^3^. Comparative analyses indicate that ECT demonstrates superior efficacy to pharmacotherapy and a faster onset of therapeutic response^4^. Beyond depression, ECT has shown clinical benefit in a range of psychiatric disorders like mania, schizophrenia, catatonia; certain uncommon indications include obsessive-compulsive disorder, neuroleptic malignant syndrome as well as specific clinical situations in Parkinson’s disease and epilepsy^4–6^. Despite its well-established efficacy, ECT remains underutilized globally, often constrained by stigma, cognitive side effects, and perceived cost-ineffectiveness compared to alternative treatments^7,8^. Utilization trends remain geographically variable, with declining rates in the United States and the United Kingdom contrasted by increasing adoption in several European and Asian countries^9–16^.

Modern ECT employs constant-current, ultra-brief, biphasic pulses (0.3–2 milliseconds (ms), 500–900 milliamperes(mA)) delivered through scalp electrodes in established montages, such as right unilateral (RUL), bitemporal (BT) or bilateral (BL), and bifrontal (BF) placements. Recently developed alternative placements, including focal electrically administered seizure therapy (FEAST) and frontomedial (FM) ECT, aim to enhance spatial targeting while minimizing cognitive adverse effects. Although the precise neurobiological mechanisms remain incompletely understood, the therapeutic efficacy of ECT is thought to depend on the magnitude and distribution of the electric field (E-field) within cortical and subcortical structures. The electrode configuration and amplitude interact with individual anatomical properties, particularly skull geometry, tissue conductivity, and cerebrospinal fluid distribution to determine the pattern and magnitude of induced current flow^17–19^. Higher stimulus amplitude is associated with improved antidepressant response but also greater cognitive side effects^20^, suggesting that the neural circuits mediating therapeutic benefit and those mediating adverse effects partially overlap but may be dissociable. E-field modeling cannot directly predict clinical outcomes, as the relationship between induced current and seizure initiation, generalization, and neuroplasticity involves numerous intermediate factors, yet it provides a spatially explicit account of which brain regions receive suprathreshold stimulation under a given electrode configuration and amplitude. If therapeutic and cognitive effects are differentially sensitive to stimulation in distinct regions, E-field distributions may offer a principled basis for comparing montages or titrating amplitude in ways that could be tested in prospective clinical studies^21^. The present modeling work is positioned within this broader translational goal.

Notably, significant interindividual and population-level differences in cranial and brain morphology have been documented. Radiographic and imaging studies report thicker frontal bones and thinner parieto-occipital bones in white males compared to black males^20^, along with differences in skull and cortical shape between Chinese and Caucasian populations^21–23^. Because the high-resistance skull is a principal determinant of current shunting during transcranial stimulation, such morphological variation is expected to alter intracerebral E-field magnitude and distribution under identical stimulation conditions, potentially influencing clinical response and seizure threshold. Consistent with a possible anatomical contribution, higher seizure thresholds have been reported among African-American^24,25^ and Japanese cohorts relative to other ethnic groups^26^. These clinical observations are difficult to interpret, however, because population differences in seizure threshold are confounded by the dose-titration procedure, concomitant medication^24^, and illness severity. Computational E-field modeling offers a complementary approach, isolating the anatomical contribution to current delivery while holding stimulation parameters constant across different modalities of stimulation. Yet systematic analyses exploring how population-level anatomical differences shape E-field magnitude and spatial distribution remain limited. Age-related brain changes increase cerebrospinal fluid (CSF)^27^. Given CSF has the highest electrical conductivity amongst all head tissues, ECT-induced E-fields may change meaningfully with age-related atrophy.

In addition to racial variation, biological sex may contribute to interindividual differences in ECT-induced electric fields. Computational modeling studies have shown that female head models generate higher cortical E-field magnitudes than male head models despite identical stimulation parameters^28–31^. These differences were associated with sex-related variation in cranial bone density and brain tissue composition, including greater proportional gray matter and white matter volumes and cortical thickness, as well as shorter scalp-to-cortex distances in females.

Our primary goal of this study was to quantitatively assess for the first time the impact of race (Chinese, Caucasian, and Black) and sex on cranial current flow (i.e. E-field magnitude) during ECT using five montages (three conventional: Bifrontal (BF), Bilateral (BL), Right Unilateral (RUL); and two experimental: Focal Electrically Administered Seizure Therapy (FEAST), and Fronto-medial (FM)). We primarily assess anatomical differences, laterality, stimulation strength, and focality across races. As a secondary analysis, we also examined the effect of race, electrode montage and sex on stimulation strength. Although it is well established that electrode montage inherently influences the spatial distribution of the induced E-field, our objective was to quantify the relative magnitude of these effects compared with those associated with age-matched and sex-balanced datasets from different cohorts.

## 2. Methods

We developed race-specific (three races: Black, Chinese, Caucasian) computational head models (N=50 (25M, 25F) per race)) to simulate ECT using five montages (three conventional and two experimental) and predicted the brain-wide E-field magnitude. Together, 750 ECT models (150 models/montage) were simulated using the Realistic vOlumetric-Approach-based Simulator for Transcranial electric stimulation (ROAST) pipeline^32^ across all subjects of different races and montages.

### 2.1 MRI

High-resolution T1-T2 weighted magnetic resonance imaging (MRI) scans were sourced for Black (1 mm^3^), Caucasian (1 mm^3^), and Chinese (1 mm^3^) race groups from the Human Connectome HCP-Young Adult 2025 project^33^ and the Chinese dataset from a single-center collection, respectively. All cohort data were age-matched and sex balanced (50% male (N=25) and 50% female (N=25); age group 20–30 yrs). For the Chinese cohort, the scanner and image acquisition parameters were given as: 48-channel radio-frequency head coil in a 3.0T GE scanner, TE: 2.96 ms, TR: 7.24 ms, flip angle: 12°, FOV: 256 x 256 mm², slice thickness = 1 mm. For Black and Caucasian groups, full imaging parameters are available at the HCP database, Appendix I: HCP Scan protocols. Briefly, structural MRI scans were acquired with Siemens MAGNETOM 3 T scanners and 32-channel head coils. T1w scan parameters were as follows: TR = 2400 ms, TE = 2.14 ms, flip angle = 8°, field of view = 224 mm x 224 mm x 180 mm, voxel size = 0.7mm^3^.

### 2.2 Montage

We considered 5 montages (3 conventional: BF, BL, RUL; and 2 experimental: FEAST, FM) as shown in Fig.1 top panel. Unless otherwise indicated, circular electrodes were modeled as disc with a 50 mm diameter (d) and 2 mm height (h), and rectangular electrodes were modeled as 63.5 mm (l) x 25.4 mm (w) x 3 mm (h). Electrode and paste compartments were simulated to mimic actual ECT administration (i.e., bottom paste layer making full scalp contact with a top electrode conductive layer). These compartments were constructed as Computer-Aided Design (CAD) files and interactively incorporated within the image data using ROAST.

1. **Bilateral (BL):** Each electrode center was positioned bilaterally at the frontotemporal positions, located 2.5 cm superior to the midpoint of the line connecting the external canthus and tragus^34^.
2. **Bifrontal (BF):** The center of each electrode was placed 5 cm superior to the lateral canthus of each eye^35^.
3. **Right Unilateral (RUL):** One electrode was placed on the right temporal scalp position (described in BL placement above) and the other electrode was placed 2.5 cm to the right of the vertex (from the edge or middle)^36^.
4. **FEAST (Focal Electrically Administered Seizure Therapy):** This montage employed two electrodes of unequal size-a small 0.75” (1.90 cm) diameter anterior circular electrode and a larger (1 x 2.5”) posterior oblong electrode. The smaller electrode’s lower boundary was positioned just above the center of the right eyebrow [16], whereas the posterior rectangular electrode was positioned tangentially to the midline with the posterior boundary 1” (2.54 cm) anterior to the vertex, and extended across the right supplementary motor cortex^37^. We note that while the sizes of the electrodes were subsequently increased for some subjects during actual administration, we still considered the first choice of dimensions in our simulations.
5. **Frontal Medial (FM):** One electrode was placed medially on the forehead and the second anterior to the vertex^36,38,39^.

**Figure 1:**
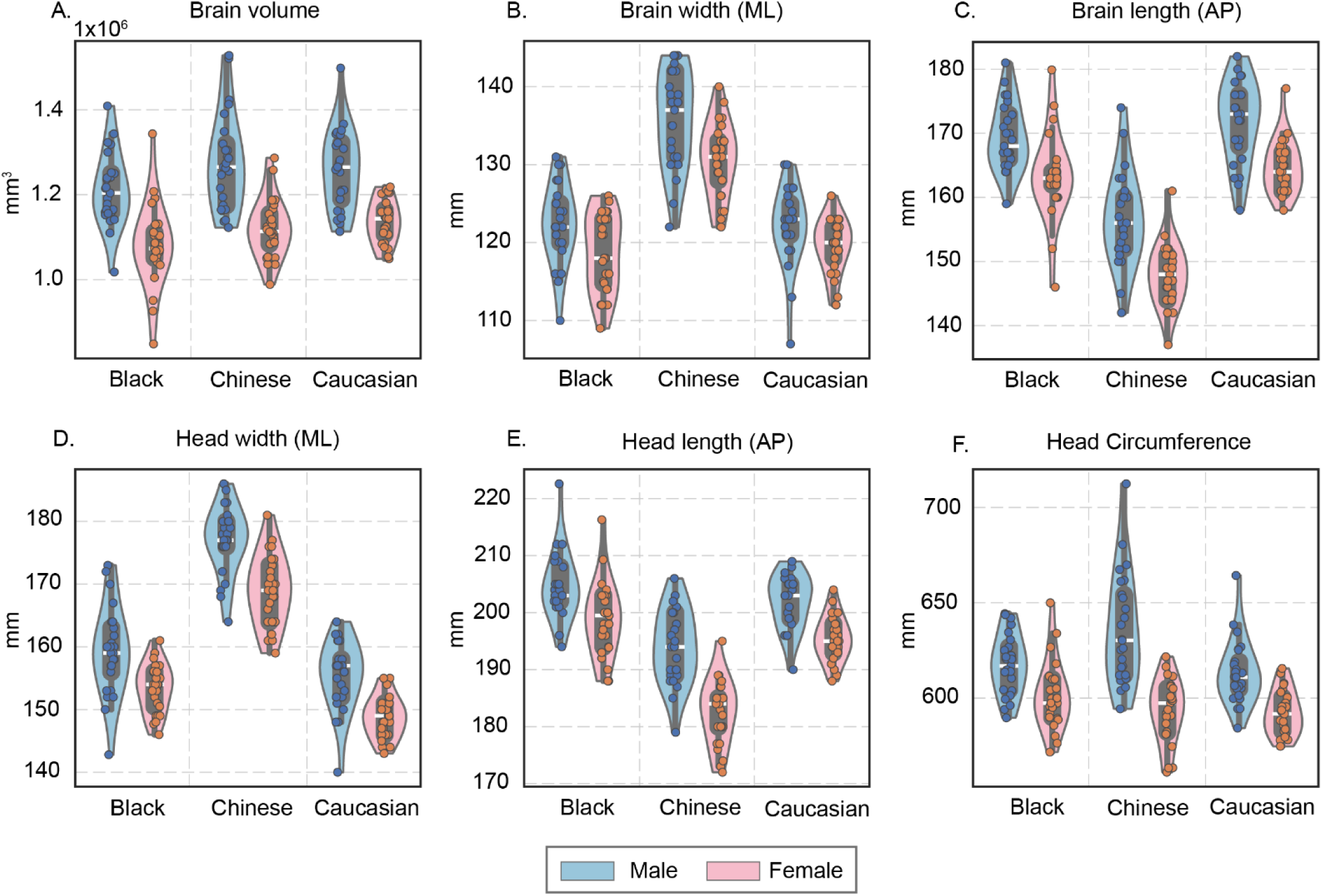
Anatomical measures (brain volume, brain width, brain length, head width, head length, and head circumference) by race and sex. Individual data points are overlaid in the violin plots, with males shown in blue and females in orange.

### 2.3 Computational Modeling

We implemented ROAST^32^ simulation workflow across the three race-specific head models and five ECT electrode montages to automate the segmentation, generate tetrahedral meshes, and solve the Laplace equation (*∇⋅`(σ`∇`V) = 0`* (*`V`*: electric potential; *∇*: gradient vector; *`σ`*: electrical conductivity) under quasi-static assumption^40,41^ to predict E-field magnitude and distribution across the brain. For each model, the MRI images, electrode locations, electrode shape and size, and conductivities were used as inputs. Briefly, the assigned isotropic electrical conductivity (S/m) values for tissues and electrode/gel compartments used in ROAST were based on our previous works^17,41^ and assigned as: skin: 0.465, fat: 0.01, skull=0.01, CSF=0.85^42^, grey matter: 0.276, and white matter: 0.126, air: 1×10^-7^, electrode: 1.4×10^6^, conductive gel: 3.5. Each simulation was solved under a constant-current boundary condition by applying 900 mA, corresponding to a clinically administered current from the anode while grounding the cathode electrode (0 mA). The interior boundaries were assigned continuity (*`n`⋅`J = 0`*; n: surface normal; *`J`* = current density) and all other external boundaries were electrically insulated (*`n`⋅`J = 0`*). Relative tolerance was set to 1×10^-6^ to improve solution accuracy. Note that all race-specific scans were resampled to 1 mm resolution prior to simulation.

### 2.4 Data analysis

Anatomical measures such as brain volume (ml), brain length (AP mm), brain width (ML mm), head length (AP mm), head width (ML mm), and head circumference (mm) were calculated across subjects for each race group (**Fig. 1**). For each montage (conventional and experimental), stimulation strength relative to neural activation threshold is calculated as *`E`_brain_`/E`_th_* metric (**Fig. 2**), where *`E`_brain_* is the 90^th^ percentile of brain-wide E-field magnitude and *`E`_th_* is E-field threshold for neuronal activation. We further calculated the ratio of median right-to-left hemisphere E-field magnitude (E) to show the laterality of predicted E-field magnitude, where ratio > 1 indicates right-dominant E-field (**Fig. 3**). Overall focality of stimulation was quantified as percentage of brain volume exposed to E-field magnitude stronger than neural activation threshold (*`E`_brain_ ≥ `E`_th_*) and robust neural activation (*`E`_brain_ ≥ `1.4*E`_th_*) (**Fig. 4**). Moreover, we assessed the relationship between stimulation amplitude and percentage of brain volume stimulated above neural activation threshold (*`E`_th_*) by linearly scaling the ECT stimulation intensity from 0 - 900 mA (with 50 mA increment) (**Fig. 5**). Finally, the effect of montage, race, and sex on stimulation strength was also computed (**Fig. 6, 7, 8, 9**).

**Figure 2:**
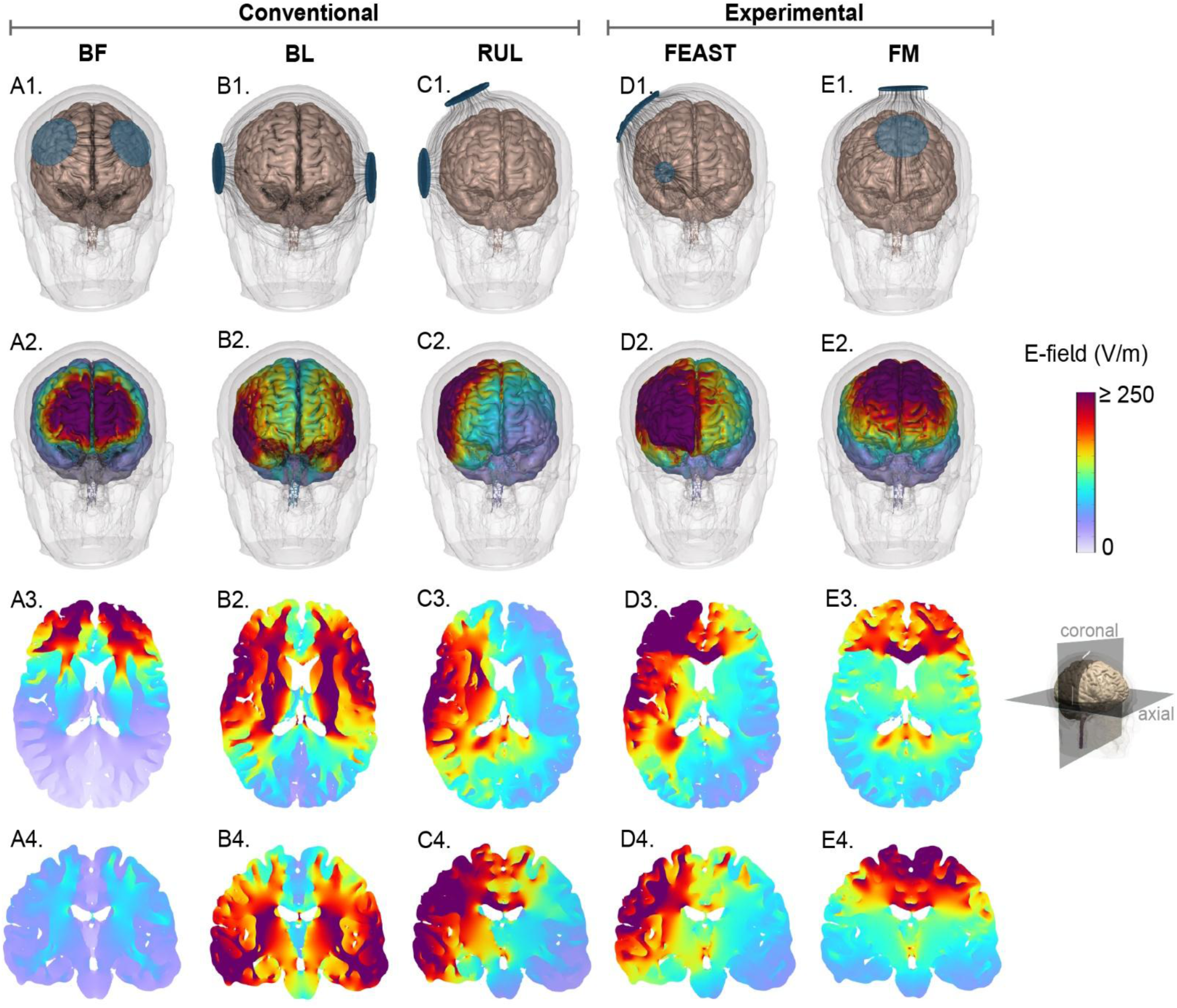
Electrode montages and corresponding E-field predictions for a representative Caucasian head model with clinically relevant ECT intensity stimulation. Panels A1-E1 depict the montages (BF, BL, RUL, FEAST, FM) with current flow streamlines across the head. Panels **A2-E2** illustrate E-field volume plot across the head, while panels A3-E3 and A4-E4 show axial and coronal slices, respectively.

**Figure 3:**
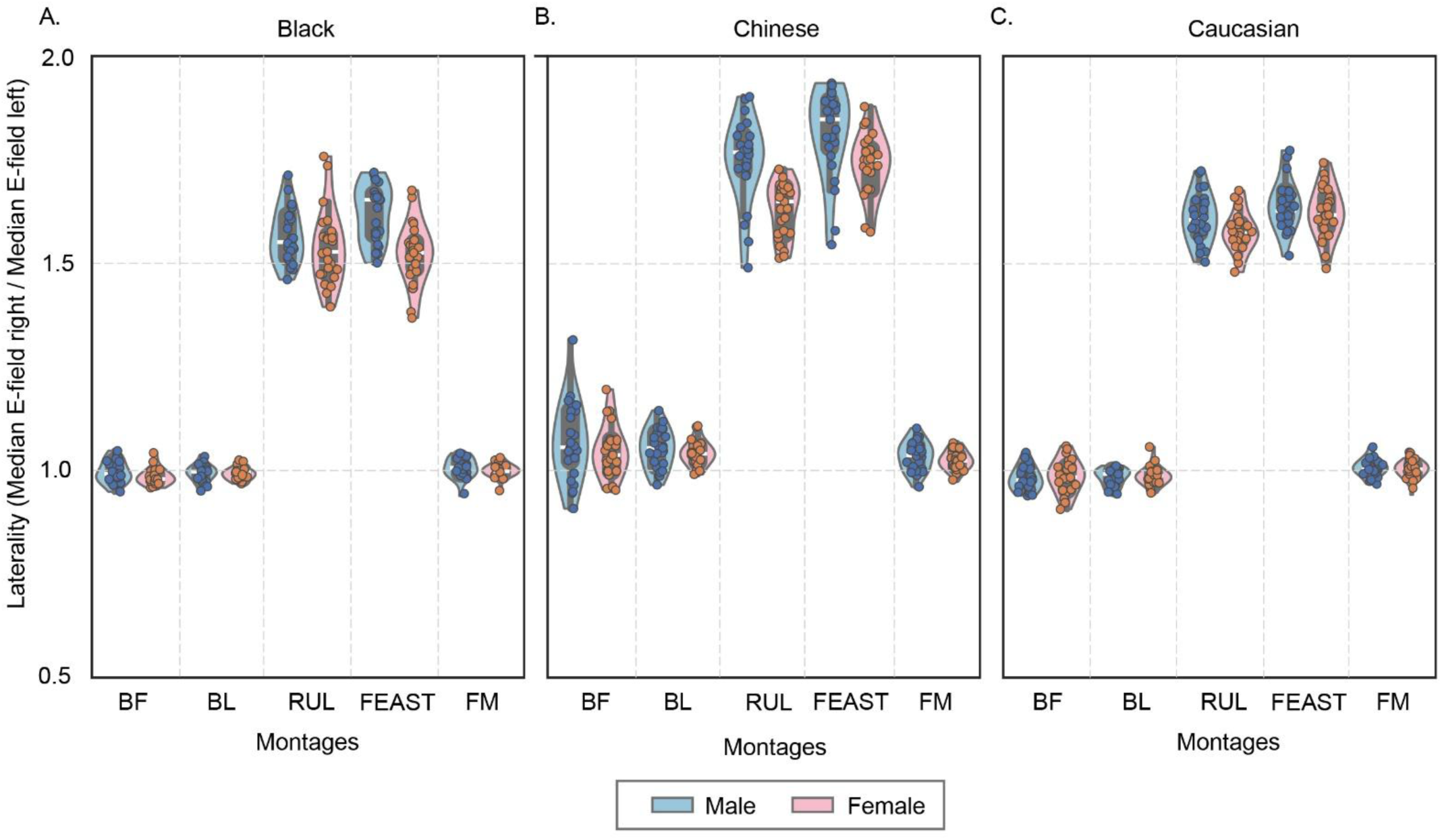
Right-to-left hemisphere median brain-wide E-field (*E*) magnitude ratio. RUL and FEAST predicted the right hemisphere dominated stimulation strength, as the laterality metric was greater than 1 (dotted line). BF, BL, and FM produced near-symmetric stimulation (laterality ratios ≈1), whereas RUL and FEAST produced substantially greater hemispheric asymmetry. Sex effects were montage dependent.

**Figure 4:**
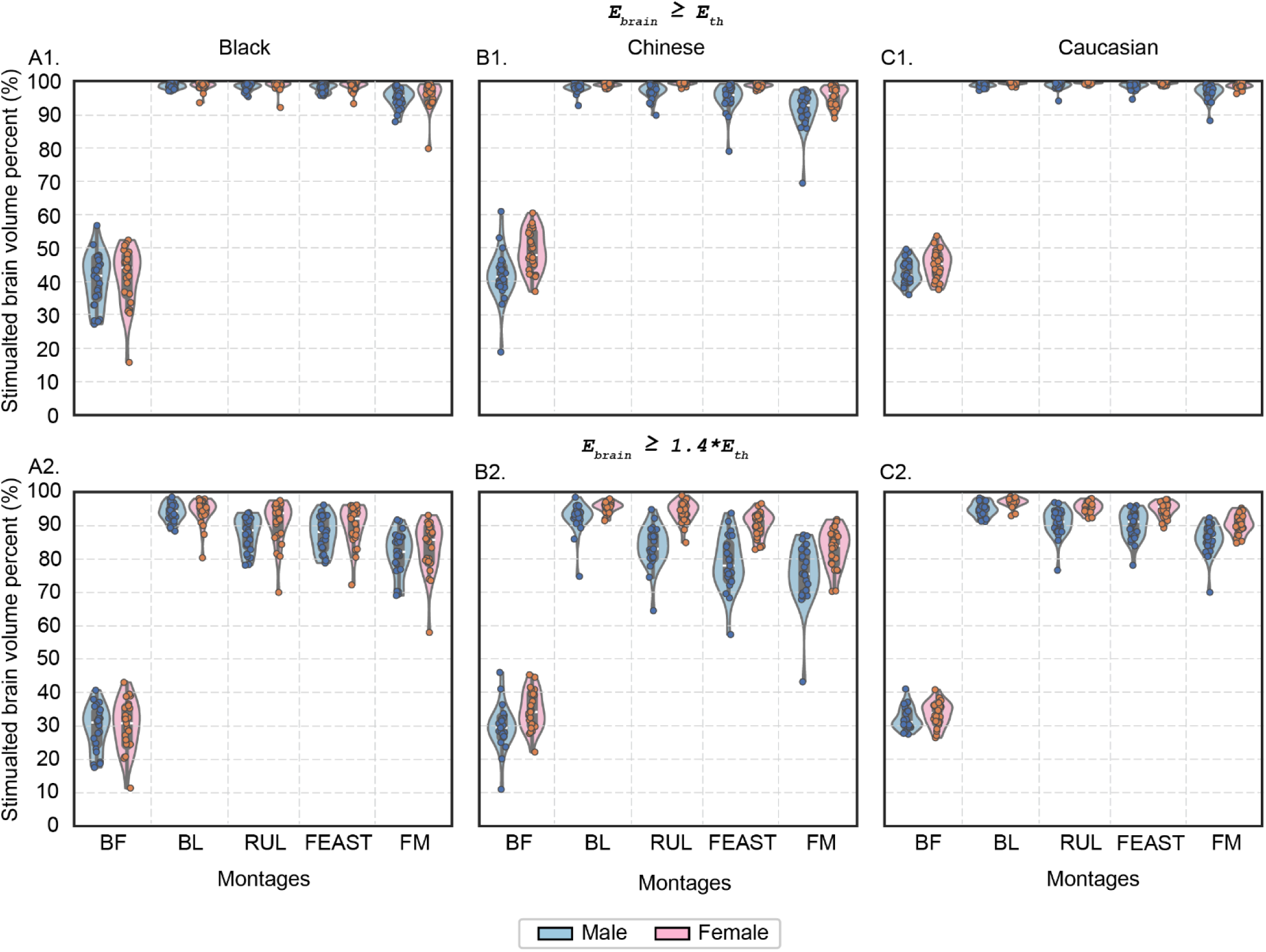
Percentage of brain volume stimulated above the neural activation threshold (*E_brain_ ≥ E_th_*, top panel) and the robust neural activation (*E_brain_ ≥ 1.4*E_th_*, bottom panel) across races and sex for conventional and experimental ECT montages. Higher percentage of *`E`_brain_ ≥ `E`_th_* and *`E`_brain_ `≥ 1.4*E`_th_* values indicate a larger proportion of brain tissue exposed to suprathreshold electric fields and therefore lower focality. Across both focality metrics, BF exhibited the highest focality, whereas BL, RUL, and FEAST exhibited the lowest focality. Caucasian subjects overall predicted lower focality than other races, while females consistently predicted lower focality than males across montages.

**Figure 5:**
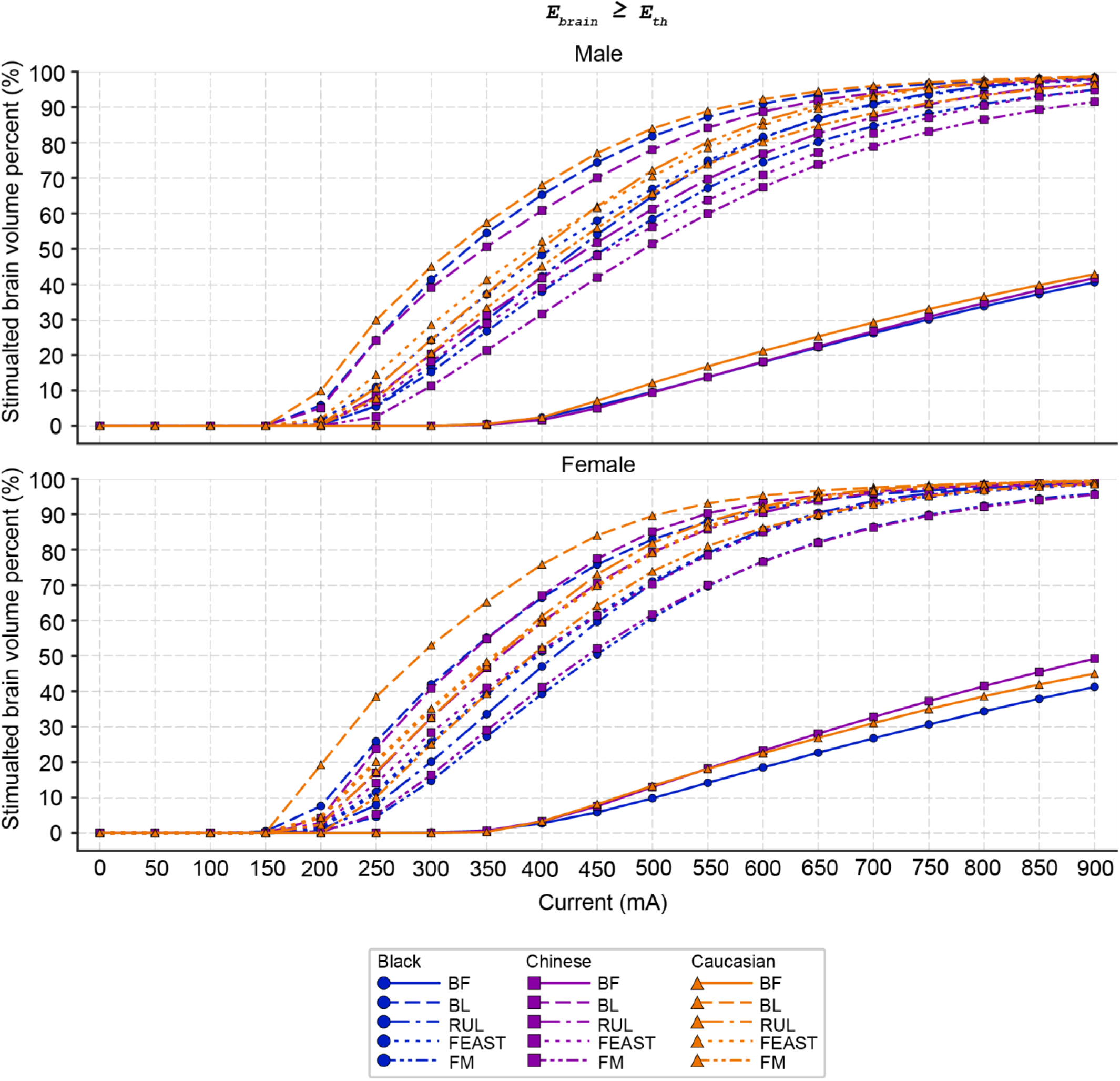
Percentage of brain volume activated above *E_th_* as a function of current intensities for each race, sex, and montage. Across all racial groups and sex, percentage of stimulated brain volume above neural activation threshold increased (therefore focality decreased) nonlinearly with ECT current intensities, exhibiting maximal focality at lower currents, rapid decreases in focality at intermediate currents, and saturation at higher current intensities. BL montage predicted the most rapid decrease in focality while the BF montage predicted the highest focality. Females predicted lower focality than males across montages.

**Figure 6:**
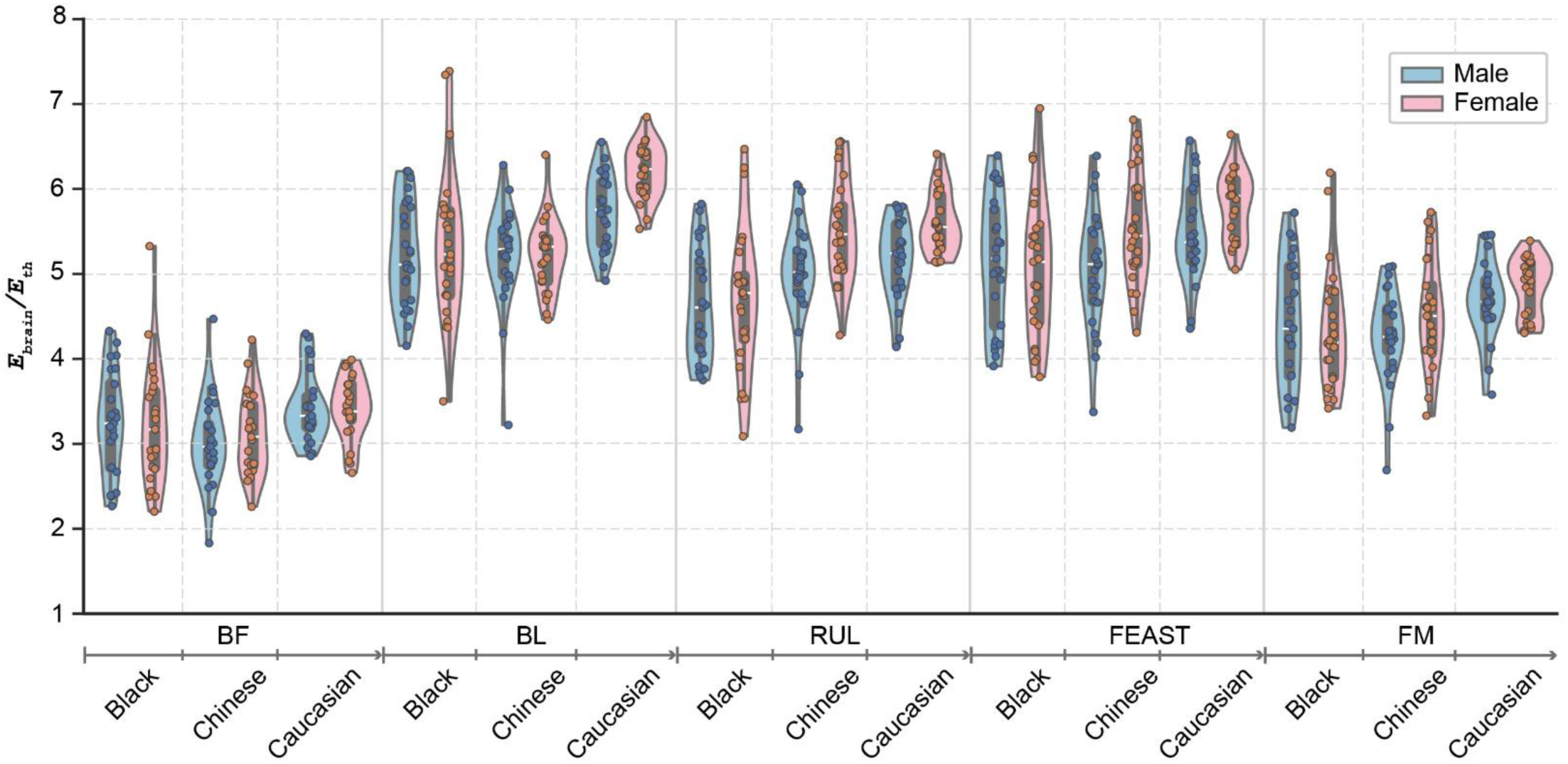
Race-related differences in stimulation strength across ECT montages. Stimulation strength varied across both montage and racial groups, with BF producing the lowest values and BL and FEAST generally producing the highest values. Race-related differences were montage dependent.

**Figure 7:**
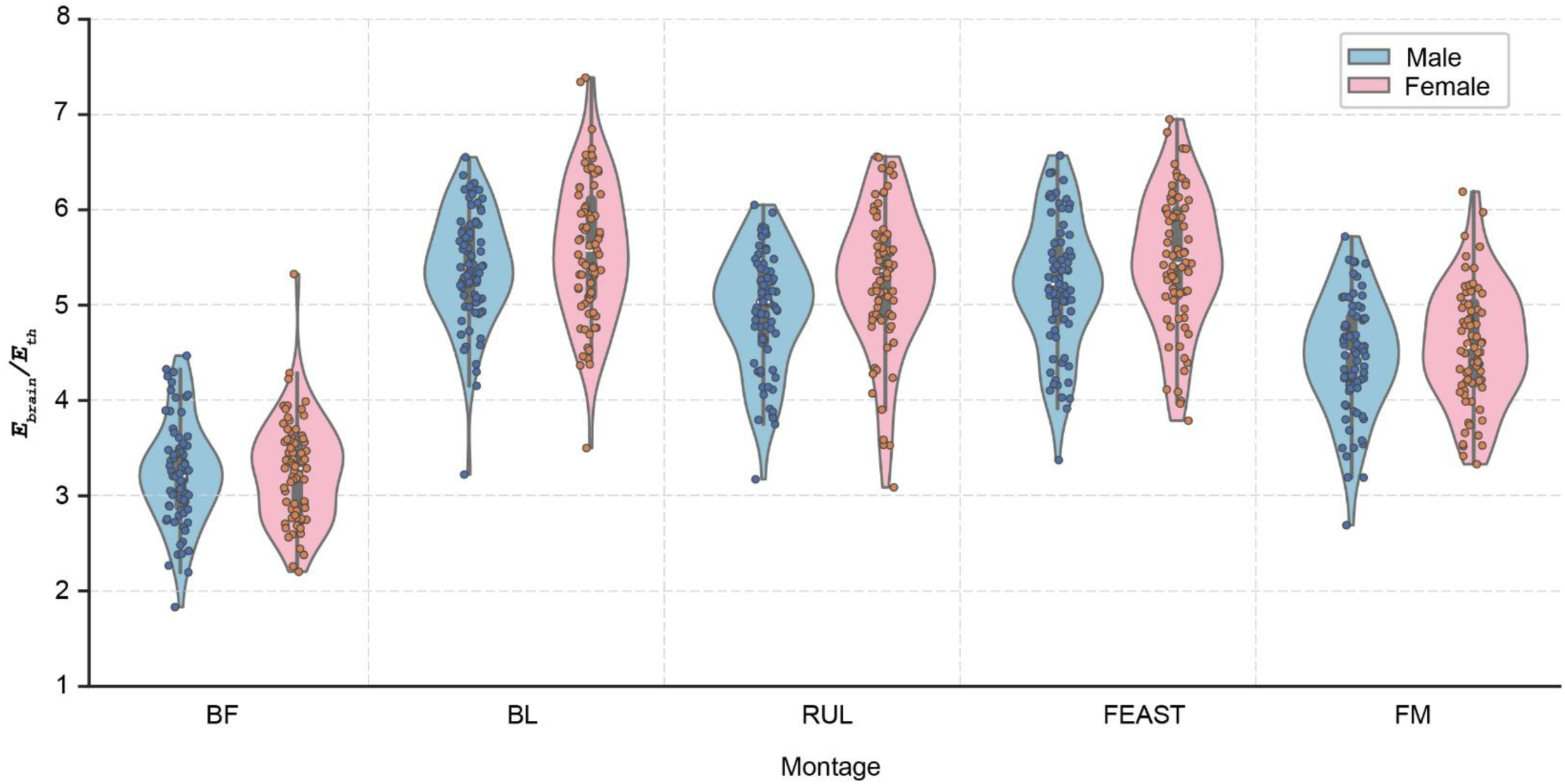
Sex-related differences in stimulation strength across ECT montages. Across montages, females generally exhibited higher stimulation strength values than males. BL and FEAST produced the highest stimulation strength, and BF produced the lowest one.

**Figure 8:**
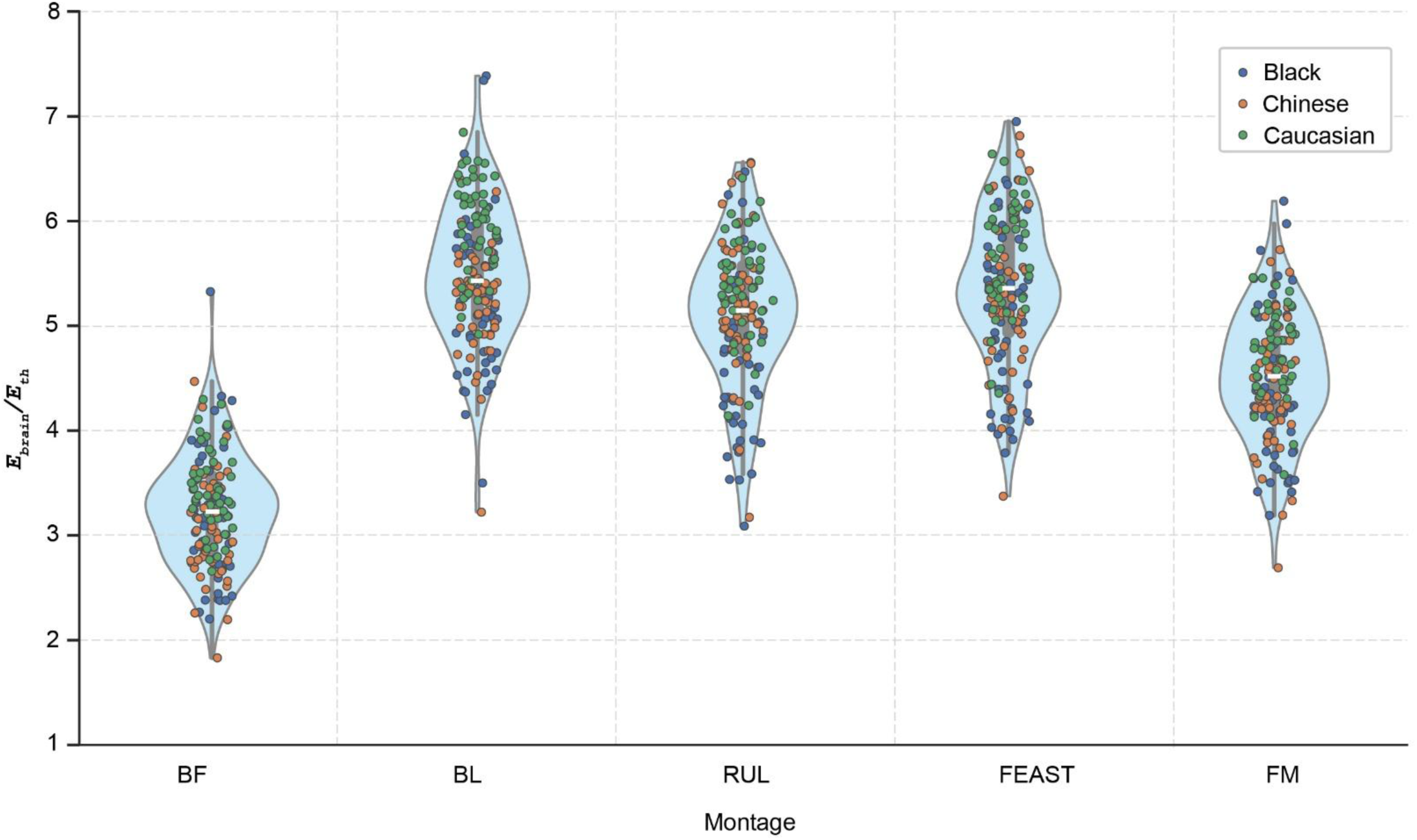
Montage-dependent differences in stimulation strength. Stimulation strength varied substantially across montage configurations, with BL and FEAST generally producing the highest stimulation strength values, BF producing the lowest values, and RUL and FM demonstrating intermediate stimulation strengths. While race-related variability was present within each montage, montage-dependent differences in stimulation strength were more pronounced than race-related differences.

### 2.4 Statistical Analysis

Statistical analyses were performed using Python (version 3.14.2)^43^ via the *statsmodels*^44^ package. Anatomical measures, including brain volume, brain width, brain length, head width, head length, and head circumference, were analyzed using two-way ANOVAs to assess the effects of race (Black, Chinese, Caucasian) and sex (male, female). For electric-field metrics (laterality stimulation strength, and overall focality), linear mixed-effects models were used to evaluate the effects of race, sex, and montage (BF, BL, RUL, FEAST, FM) while accounting for repeated measurements within subjects. When significant interactions were identified, post hoc pairwise comparisons were performed using Tukey’s honestly significant difference (HSD) procedure, followed by FDR correction. To evaluate the homogeneity-of-variance assumption underlying the linear mixed-effects models, Brown-Forsythe tests were applied to model residuals. Statistical significance was defined as *`α < 0.05`*.

## 3. Results

### 3.1 Anatomical measures

Race was associated with several anatomical characteristics, including brain volume (*F* (2,144) =5.98, *p*=0.003, η²ₚ=0.077), brain width (*F* (2,144) =55.93, *p*<0.001, η²ₚ=0.437), brain length (*F* (2,144) =113.32, *p*<0.001, η²ₚ=0.612), head width (*F* (2,144) =225.90, *p*<0.001, η²ₚ=0.758), and head length (F (2,144) =79.39, *p*<0.001, η²ₚ=0.524), whereas head circumference did not differ significantly among racial groups after FDR correction (*F* (2,144) =2.49, *p*=0.087). Sex influenced all anatomical measures (all FDR-adjusted *p≤0.001*), with males exhibiting larger brain and head dimensions than females. No race and sex interactions were detected for brain volume, brain width, brain length, head width, or head length (all FDR-adjusted *p*>0.05). Head circumference only exhibited a significant race and sex interaction (*F* (2,144) =6.08, FDR-adjusted *p*=0.018, η²ₚ=0.078). A follow-up post-hoc analyses showed racial differences among males, with Chinese males exhibiting larger head circumferences than both Black (FDR-adjusted *p*=0.011) and Caucasian males (FDR-adjusted *p*=0.003). No racial differences were identified among females (all FDR-adjusted *p≥*0.34) (**Fig. 1**).

### 3.2 E-field spatial profile and intensity

**Fig. 2** shows an exemplary Caucasian head model with the five montages considered in the study, together with the corresponding E-field magnitude plots (3D and cross-sectional slice). Prediction shows BL montage with the most extensive coverage in both hemispheres. RUL and FEAST predicted more spatially constrained patterns and hemispheric asymmetric patterns whereas FM predicted concentrated E-fields within superior frontal regions. BF montage predicted the most localized and weakest fields compared to the other montages.

Unless otherwise stated, most of our analyses were based on 90^th^ percentile of brain-wide E-field magnitude (*`E`_brain_*) and the *`E`_brain_* magnitudes for each montage, racial group, and their corresponding sex are summarized in Table 1. Across all racial groups, BL montage predicted the highest *`E`_brain_* magnitudes, followed by FEAST and RUL, whereas BF consistently produced the lowest E-field magnitudes. Caucasian heads generally predicted the highest *`E`_brain_* magnitudes whereas the Black heads predicted the lowest values. Female heads tended to predict slightly higher *`E`_brain_* magnitudes than males, particularly for BL, FEAST, and RUL montages.

**Table 1:** Brain-wide E-field magnitude (*E_brain_*) across montages for different races and the corresponding sex. Values are reported as mean, SD, and range.

| Race | E-field magnitude ($E_{brain}$, V/m) | | | | | |
| --- | --- | --- | --- | --- | --- | --- |
|  | Sex | Montages |  |  |  |  |
|  |  | BF | BL | RUL | FEAST | FM |
| Black | Male | 81.41 ± 14.89<br>(56.69 -108.1) | 130.90 ± 15.25<br>(103.83 -155.2) | 116.12 ± 16.38<br>(93.76 -145.60) | 128.52 ± 19.20<br>(97.87-159.84) | 110.59 ± 18.28<br>(79.77-143.00) |
|  | Female | 80.28 ± 17.48<br>(55.06 -133.1) | 133.68 ± 22.27<br>(87.52 -184.66) | 117.62 ± 21.29<br>(77.20 -161.74) | 127.07 ± 20.69<br>(94.69 -173.74) | 108.77 ± 17.80<br>(85.44 -154.76) |
| Chinese | Male | 75.19 ± 13.11<br>(45.78 -111.73) | 130.44 ± 14.55<br>(80.55 -157.01) | 124.83 ± 15.14<br>(79.31 -151.28) | 126.15 ± 16.99<br>(84.35 -159.73) | 106.87 ± 13.87<br>(67.27 -127.24) |
|  | Female | 77.80 ± 11.82<br>(56.48 -105.6) | 130.64 ± 10.64<br>(111.56 -160.01) | 137.76 ± 14.93<br>(106.97 -163.9) | 138.03 ± 16.42<br>(107.74 -170.36) | 113.57 ± 15.87<br>(83.26 -143.15) |
| **Caucasian** | Male | 85.73 ± 10.19<br>(71.43 -107.4) | 143.19 ± 10.72<br>(123.01 -163.8) | 129.35 ± 11.55<br>(103.52 -145.2) | 137.18 ± 13.99<br>(108.94 -164.25) | 117.61 ± 11.75<br>(89.48 -136.51) |
|  | Female | 85.36 ± 9.23<br>(66.47- 99.74) | 155.02 ± 7.93<br>(138.27-171.1) | 139.90 ± 8.99<br>(128.29 -160.2) | 144.88 ± 10.04<br>(126.28 -166.00) | 121.32 ± 8.50<br>(107.64 -134.7) |

As shown in **Table 1**, variability in *`E`_brain_* differed across race-sex groups. Brown-Forsythe test of variance homogeneity conducted on the raw *`E`_brain_* values showed significant differences in variance across race-sex groups for BL, RUL, FEAST, and FM (all *p*<0.05), but not for BF (*p*=0.0806). However, the tests on the residuals from the linear mixed-effects models showed no significant heterogeneity of variance for all montages (all *p*≥0.28), indicating no evidence of residual heteroscedasticity. These findings indicate that although variability differed across demographic groups in the raw E-field data, residual variance remained comparable across groups after accounting for race, sex, and montage, and repeated measurements within the linear mixed-effects model.

### 3.3 Laterality of stimulation

Laterality was influenced by race (*F* (2,720) =196.1, *p*<0.001), sex (*F* (1,720) =51.3, *p*<0.001), and montage (*F* (4,720) =5150.8, *p*<0.001), with significant race and sex, race and montage, sex and montage, and three-way interactions (all *p*<0.01). Across racial groups, laterality ratios were greater in Chinese subjects than in both Black (mean difference=0.098, FDR-adjusted *p*=0.0019) and Caucasian subjects (mean difference=0.078, FDR-adjusted *p*=0.0119), whereas Black and Caucasian subjects did not differ significantly (FDR-adjusted *p*=0.465). Males exhibited greater laterality ratios than females during RUL and FEAST stimulation (mean differences=0.060 and 0.064, respectively; FDR-adjusted *p*=0.0007 and 0.0019), whereas no sex differences were observed for BF, BL, or FM stimulation (all FDR-adjusted *p≥0.137*. Across montage, BF, BL, and FM predicted comparable laterality ratios (all FDR-adjusted *p*>0.86), whereas RUL and FEAST produced substantially greater laterality ratios than the remaining montages (all FDR-adjusted *p*<0.001). FEAST further exceeded RUL (mean difference=0.0495, FDR-adjusted *p*<0.001), indicating the strongest hemispheric asymmetry among the montages evaluated (**Fig. 3**).

### 3.4 Overall focality at a clinically relevant intensity (comparison of neural and robust neural activation threshold)

Focality was quantified as the percentage of stimulated brain volume exceeding the neural activation threshold (*`E`_brain_ `≥ E`_th_*) and the robust neural activation threshold (*`E`_brain_ `≥ 1.4*E`_th_*). A higher percentage of *`E`_brain_ `≥ E`_th_* and *`E`_brain_ `≥ 1.4*E`_th_* depict less focal montage or larger brain volume exposed over the neural activation threshold. For *`E`_brain_ ≥ `E`_th_*, there was a significant effects of race (*F* (2,720) =11.66, *p*<0.001), sex (*F* (1,720) =52.65, *p*<0.001), and montage (*F* (4,720) =7143.61, *p*<0.001), as well as race and sex (*F* (2,720) =13.30, *p*<0.001), race and montage (*F* (8,720) =8.14, *p*<0.001), and sex and montage (*F* (4,720) =3.70, *p*=0.005) interactions. The three-way interaction was not significant (*F* (8,720) =1.52, *p*=0.147).

Race-related differences varied across montage configurations. For BF montage, Chinese subjects exhibited lower focality than Black subjects (mean difference=0.046, FDR-adjusted *p*=0.009), whereas focality in Caucasian subjects did not differ significantly from either group. For BL and RUL stimulation, Caucasian subjects exhibited lower focality than Chinese subjects (mean differences=0.007 and 0.011, respectively; both FDR-adjusted *p*=0.005). For FEAST stimulation, both Caucasian and Black subjects exhibited lower focality than Chinese subjects (mean differences=0.022 and 0.015, respectively; both FDR-adjusted *p*≤0.009). For FM stimulation, Caucasian subjects exhibited lower focality than both Chinese (mean difference=0.040, FDR-adjusted *p*<0.001) and Black subjects (mean difference=0.021, FDR-adjusted *p*=0.032). Females showed lower focality than males across all montages (mean differences=0.005-0.035; all FDR-adjusted *p*≤0.003). Montage-wise, BF produced higher focality than BL, RUL, FEAST, and FM (mean differences=0.552, 0.551, 0.545, and 0.521, respectively; all FDR-adjusted *p*<0.001). BL, RUL, and FEAST focality did not differ significantly and were lower than FM (mean differences=0.031, 0.030, and 0.024, respectively; all FDR-adjusted *p*<0.001) (**Fig. 4**).

A similar pattern was observed for *`E`_brain_ ≥ `1.4*E`_th_*. Significant effects of race (F (2,720) =56.96, *p*<0.001), sex (F (1,720) =125.21, *p*<0.001), and montage (F (4,720) =3816.98, *p*<0.001) were identified, together with race and sex (F (2,720) =21.20, *p*<0.001), race and montage (F (8,720) =6.75, *p*<0.001), and sex and montage (F (4,720) =6.10, *p*<0.001) interactions. The three-way interaction was not significant (F (8,720) =0.94, p=0.486). For BF stimulation, Caucasian subjects exhibited lower focality than Black subjects (mean difference=0.033, FDR-adjusted *p*=0.027). For BL stimulation, Caucasian subjects did not exceed in focality than both Black (mean difference=0.017, FDR-adjusted *p*=0.027) and Chinese subjects (mean difference=0.019, FDR-adjusted *p*=0.017). A similar pattern was observed during RUL stimulation, where Caucasian subjects exhibited lower focality than both Black (mean difference=0.040, FDR-adjusted *p*=0.008) and Chinese subjects (mean difference=0.043, FDR-adjusted *p*=0.004). For FEAST stimulation, Caucasian subjects exhibited lower focality than Chinese subjects (mean difference=0.079, FDR-adjusted *p*<0.001), while Black subjects likewise did not exceed Chinese subjects (mean difference=0.043, FDR-adjusted *p*=0.008). For FM stimulation, Caucasian subjects exhibited lower focality than both Black (mean difference=0.057, FDR-adjusted *p*=0.001) and Chinese subjects (mean difference=0.087, FDR-adjusted *p*<0.001). Females consistently showed lower focality than males across all montages (mean differences=0.018 - 0.063; all FDR-adjusted *p*≤0.021). BL produced lower focality than both RUL and FEAST (mean differences=0.042 and 0.060, respectively; both FDR-adjusted *p*<0.001), whereas RUL and FEAST did not differ significantly (**Fig. 4**).

### 3.5. Stimulation focality across ECT intensities

The percentage of stimulated brain volume exceeding the neural activation threshold across ECT current intensities (0 - 900 mA with a 50 mA increment) is shown in **Fig. 5**. Higher focality refers to a lower percentage of stimulated brain volume. Linear scaling from the 900 mA baseline was applied due to linearity in the E-field distribution. A non-linear relationship characterized current amplitude and focality (*`E`_brain_ ≥ `E`_th_*), with higher focality at lower current intensities (<200-250 mA), rapid decreases at intermediate currents, and saturation at higher currents. Montage-dependent differences were substantially larger than race- or sex-related differences. BL montage produced the most rapid decrease in focality, reaching near-complete stimulation of brain volume (>95%) by approximately 700–900 mA in all racial groups, whereas BF produced only ∼40%–50% of stimulated brain volume percentage even at 900 mA. RUL, FEAST, and FM demonstrated intermediate focality profiles, with group differences most evident within the steeply rising portion of the stimulated brain volume percentage curves. Caucasian subjects generally exhibited lower focality for BL, RUL, FEAST, and FM montages, whereas Chinese subjects predicted the highest focality. Females exhibited lower focality than males across all montages, with the largest deviations noted in the RUL, FEAST, and FM montages. As ECT intensity approached saturation, differences between races became less pronounced.

### 3.6 Effect of Race

Race significantly influenced stimulation strength (*F* (2,720) =48.96, *p*<0.001), and this effect was montage dependent (*F* (8,720) =4.25, *p*<0.001). Across montages, Caucasian subjects generally exhibited higher stimulation strength (*`E`_brain_* `/E`_th_). For BF montage, Caucasian subjects predicted higher stimulation strength compared to Chinese (mean difference=0.362, FDR-adjusted *p*=0.0048) subjects. Caucasian subjects exceeded *`E`_brain_`/E`_th_* than both Black and Chinese subjects for BL montage (mean differences=0.673 and 0.743, respectively; both FDR-adjusted *p*<0.001). For RUL montage, both Caucasian and Chinese subjects exhibited greater *`E`_brain_`/E`_th_* values than Black subjects (mean differences=0.710 and 0.577, respectively; both FDR-adjusted *p*<0.001), whereas Caucasian and Chinese subjects did not differ significantly. A similar pattern was observed for FEAST and FM stimulation, where Caucasian subjects exhibited higher *`E`_brain_`/E`_th_* than both Black and Chinese subjects (FEAST: mean differences = 0.529 and 0.358; FDR-adjusted *p*=0.001 and 0.041, respectively; FM: mean differences = 0.392 and 0.370; FDR-adjusted *p*=0.007 and 0.011, respectively) (**Fig. 6**).

### 3.7. Effect of Sex

Sex differences also contributed to variability in stimulation strength (*F* (1,720) =17.30, *p*<0.001). Across all five montages, females exhibited modestly higher *`E`_brain_`/E`_th_* values than males, including BF (3.25 vs 3.23), BL (5.59 vs 5.39), RUL (5.27 vs 4.94), FEAST (5.47 vs 5.22), and FM (4.58 vs 4.47). However, the absence of a significant sex and montage interaction (*F* (4,720) =1.57, *p*=0.180) indicates that this difference was relatively consistent across stimulation montages, suggesting that sex influences overall *`E`_brain_`/E`_th_* without substantially altering montage-dependent predictions (**Fig. 7**).

### 3.8 Effect of Montage

Montage was the dominant determinant, with a strong effect on *`E`_brain_`/E`_th_* values (*F* (4,720) =358.39, *p*<0.001). BF montage predicted substantially lower *`E`_brain_`/E`_th_* than BL, FEAST, RUL, and FM (mean differences=1.286 - 2.254; all FDR-adjusted *p*<0.001). BL and FEAST montages predicted comparable *`E`_brain_`/E`_th_* (mean difference=0.147, *p*=0.287), and both montages exceeded RUL (mean differences=0.389 and 0.242, respectively) and FM (mean differences=0.968 and 0.821, respectively). RUL montage also exceeded *`E`_brain_`/E`_th_* than the one predicted by FM montage (mean difference=0.579, FDR-adjusted *p*<0.001). There was no sex and montage interaction (*F* (4,720) =1.57, *p*=0.180), indicating a similar montage-dependent prediction pattern in males and females (**Fig. 8**).

## 4. Discussion

In this modeling study, we investigated how race-, sex-, and anatomy-related variability influence the ECT-induced E-field across five conventional and experimental ECT montages, namely BF, BL, RUL, FEAST, and FM, respectively. Three key findings emerged. First, montage was the dominant determinant of stimulation outcomes, exerting substantially larger effects than either race or sex across tested metrics (stimulation strength, focality, and laterality). Second, race-related anatomical differences led to systematic changes in predicted E-field metrics, particularly in stimulation strength and focality. Third, although sex-related effects were generally smaller, females consistently exhibited higher stimulation strengths but lower focality than males.

### 4.1 Anatomical differences vary across race and sex

Substantial anatomical variation was observed across racial groups. Chinese subjects exhibited markedly greater left-right brain and head dimensions but smaller anterior-posterior dimensions relative to Black and Caucasian subjects. These findings highlight that cranial anatomy varies in a multidimensional way and cannot be adequately characterized by a single measure, such as intracranial volume or head circumference alone, consistent with previous neuroimaging studies demonstrating that brain morphology reflects complex regional and global structural variation rather than simple differences in overall size^45,46^. Indeed, head circumference did not differ significantly among racial groups despite pronounced differences in brain and head shape. This suggests that cranial geometry, rather than overall head size alone, may be an important determinant of E-field measures, consistent with prior modeling studies showing that interindividual anatomical variability substantially alters ECT stimulation strength and focality^47^.

Sex-related differences were observed across all anatomical measures, with males exhibiting larger brain and head dimensions than females, consistent with prior literature^45,46^. However, larger anatomy did not translate into stronger stimulation. Instead, females generally demonstrated greater stimulation strength and a higher percentage of stimulated brain volume relative to neural activation threshold (less focality). This observation suggests that anatomical scaling relationships influencing E-field propagation may be more complex than simple differences in head size and likely involve interactions among head geometry, brain dimensions, and montage relative to the underlying cortex^47^.

### 4.2 Montage selection governs stimulation strength

Montage configuration exerted the strongest influence on stimulation strength (*`E`*_brain_/*E*_th_). Across racial groups, BL and FEAST consistently exhibited the greatest stimulation strengths, followed by RUL and FM, whereas BF generated the lowest values. The magnitude of montage-related differences was substantially greater than any race- or sex-related effect, underscoring electrode placement as the principal determinant of ECT dose delivery. Previous modeling studies have shown that electrode montage and current-flow pathways strongly influence E-field distribution^19,47,48^. Consistent with this biophysical principle, substantial montage-dependent differences in E-field outcomes were observed in the present study as BL and FEAST montages concentrate current flow through large portions of the cerebral hemispheres, resulting in greater field magnitudes throughout the brain. In contrast, BF produced lower stimulation strengths despite activating broad frontal regions, indicating that E-field intensity and stimulated volume represent distinct aspects of ECT dose distribution^19,34^.

### 4.3 Race contributes to variability in ECT outcomes

Race significantly influenced stimulation strength, focality, and laterality. However, these effects were highly montage dependent. Caucasian subjects generally exhibited the highest stimulation strengths across BL^49^, FEAST, FM, and focality metrics. In contrast, Chinese subjects exhibited the greatest hemispheric laterality. The focality analyses further demonstrated that racial effects depended strongly on electrode configuration. For most montages, Caucasian subjects exhibited lower focality than Black and Chinese subjects. The most notable exception was BF stimulation, where Chinese subjects exhibited lower focality than Black subjects. Interestingly, racial differences were most evident under conditions where montage-dependent differences were already pronounced. This indicates that demographic anatomy does not simply shift all E-field metrics uniformly; rather it interacts with the current-flow pattern imposed by a specific electrode montage. Consequently, patient-specific anatomical factors may become particularly important when selecting montages designed to optimize stimulation outcomes.

### 4.4 Sex influences stimulation efficiency

Although sex-related effects were smaller than montage effects, they were remarkably consistent across analyses. Females exhibited higher stimulation strengths but lower focality across nearly all montages. These findings are largely attributed to males possessing larger brain and cranial dimensions^27^. One possible interpretation is that larger cranial dimensions increase current dispersion before reaching cortical targets, reducing normalized stimulation efficiency^53^. While the present study was not designed to isolate the mechanistic contribution of individual anatomical factors, the consistency of the observed sex-related effects suggests that sex-specific anatomical characteristics systematically alter ECT dose delivery. Notably, sex-related effects did not substantially alter the relative ranking of montages. The absence of a significant sex and montage interaction for stimulation strength indicates that montage-dependent patterns were preserved across sex. Thus, sex contributes to overall stimulation efficiency without fundamentally changing the comparative behavior of the different electrode montages.

### 4.5 Distinct montage effects on laterality and focality

Laterality analyses showed a pattern distinct from stimulation strength and focality. FEAST produced the highest laterality ratios, followed by RUL, whereas BF, BL, and FM demonstrated near-symmetric stimulation. These findings confirm that FEAST and RUL preferentially drive hemispherically asymmetric current flow, whereas BF and BL distribute current more bilaterally.

The focality analysis demonstrated that BF consistently stimulated smaller percentages of brain volume relative to neural activation threshold (more focal) than BL, RUL, and FEAST montages, with BL generally stimulating a higher percentage of brain volumes above neural activation threshold (less focal). Thus, a trade-off was apparent across montages. BF maximized focality and but minimized stimulation strength, FEAST maximized hemispheric asymmetry, and BL maximized stimulation strength but minimized focality. The focality vs ECT intensity analyses further demonstrated that montage-dependent differences were preserved across a broad range of stimulation amplitudes.

### 4.6 Clinical Implications

Our findings suggest that ECT outcome is influenced by both electrode montage and patient anatomy. Montage selection accounted for the largest proportion of variability in predicted E-field measures and therefore remains the most important factor for clinical trial design. However, race- and sex-related anatomical variation produced measurable differences in stimulation strength, focality, and laterality that were comparable in magnitude to some montage-specific contrasts. These findings support use of patient-specific anatomy when individualized ECT dosing strategies based on E-field metrics are considered. Computational modeling has traditionally focused on montage optimization, but the current results demonstrate that demographic anatomy can systematically bias stimulation outcomes even when identical stimulation parameters are applied.

### 4.7 Model validity, limitations and future directions

In this study, BF montage consistently produced the lowest overall cortical E-field magnitude across the cohort, reflecting increased scalp shunting between closely spaced anterior electrodes (50 mm diameter)^54^. Moreover, the ECT E-field prediction for the BF montage is consistent with a prior ECT modeling paper^55^. Unlike prior ECT modeling studies^56^ that employed truncated head models, our pipeline automatically segmented tissue masks below the truncated boundary^32^ to improve modeling results^57^. Our study leveraged individualized head models across a large cohort, enabling characterization of population-level anatomical variability, whereas many prior modeling studies have relied on a single representative head model^56,58,59^.

Several limitations of this study should be acknowledged. The analyses were based on computational head models rather than direct physiological or clinical outcome measures. Consequently, the observed differences represent predicted alterations in E-field distribution rather than confirmed differences in seizure induction or treatment response. The age group selected for this study (20-30 years old) is based on the HCP-Young Adult 2025 database, which only includes ages 22-35 years and is considered relatively younger than a typical ECT population^60–63^. Therefore, the generalizability of the findings may be limited. Race was used as a grouping variable to characterize anatomical variability and should not be interpreted as a biological determinant of treatment outcome. The observed effects are mediated by underlying anatomical differences represented within the head models. Although the sample included multiple racial and sex groups, additional populations and age-related anatomical variability were not examined. Our modeling approach does not account for factors such as differences in susceptibility to E-fields including seizure threshold (e.g, charge / number of pulses), across brain regions. Consequently, clinical outcomes cannot be inferred directly from these simulations and must be established empirically. The scans for Black and Caucasian subjects were sourced from the HCP database (pooled from multiple centers) and Chinese subjects from a single center, which may introduce notable technical differences in preprocessing pipelines and thus impact downstream analysis. We did not explicitly model a fat tissue compartment whose resistive property can minimize shunting and more current penetration through the scalp^64–66^. Therefore, inclusion of fat compartment is not expected to alter the relative current flow pattern and the overall results. Future studies should investigate whether the demographic differences identified here translate into measurable differences in seizure expression, cognitive side effects, or antidepressant response. Integration of individualized head modeling into clinical ECT planning may provide a pathway toward precision-guided stimulation strategies tailored to patient-specific anatomy.

### 4.8 Conclusions

Montage was the dominant determinant of predicted ECT E-field measures, exerting substantially larger effects than race or sex across metrics of stimulation strength, focality, and laterality. BL and FEAST generated the highest stimulation strengths, FEAST produced the greatest hemispheric asymmetry, and BF produced the highest focality (lower stimulated brain volume percentage relative to the neural activation threshold). Race-related anatomical variation systematically altered stimulation strength and focality, with Caucasian head models generally exhibiting lower focality and stimulation strength, whereas Chinese head models demonstrated the greatest laterality. Females consistently exhibited higher stimulation strengths but lower focality (larger activated brain volumes) than males. Together, these findings demonstrate that demographic anatomy contributes to variability in ECT outcomes and supports the incorporation of patient-specific anatomical information into future ECT optimization and treatment-planning.

## Acknowledgements

Sources of financial support: This study was partially funded by grants to NK, AD, CA from NIH. NK is supported by R01DA060914, 1R21NS136909-01. AD is supported by 1UG3DA063344-01, R01DA060914, and 1UG3NS139014-01. CCA is supported by MH128692 and MH125126.

## Data availability statement

All data that supports the findings of this study are included within the article (and any supplementary files).

